# BIOCHEMICAL COMPOSITION OF SPECIALTY TEA EXTRACTS FROM MULTISTAGE EXTRACTIONS USING SOLVENTS TREATMENTS

**DOI:** 10.1101/2025.01.29.635473

**Authors:** Emily Kipsura, Robert Koech, Simon Mwangi, Samson Kamunya, Victor Ngeno, Joel Koech, Christine Bii

## Abstract

Diversification and value addition in tea has received considerable attention in recent past. This has been occasioned by increased tea volumes in the world market, especially the aerated product that has led to over flooding. Additionally, health benefits associated with diversified products such as green tea and extracts has elicited a lot of interests in both the industry and external players in finding out alternative ways of tea consumption. The aim of this study was to determine biochemical composition of the various tea extract products generated at different points of a multistage extraction process. The extraction process consisted of hot distilled water, methylene chloride and ethyl acetate steps. The products of purple, yellow, green, black orthodox teas, matcha and the semi aerated oolong were used as raw materials for the extracts. Total polyphenols content, individual catechins and caffeine content were assayed using UV-spectrophotometer and High Performance Liquid Chromatography-Photo diode array HPLC-PDA. Analysis of variance (ANOVA) was used for statistical purposes to detect significant differences between the extraction methods with the least significant difference (LSD) used to separate the means in the assayed tea samples. Results generated indicated that total polyphenols, individual catechins and caffeine content were significantly different (p≤0.05) in the assayed tea samples. Water extracts showed high total yield (82.2%), while methylene chloride recorded the least (2.63%). Ethyl acetate tea extracts showed high phenolic content that ranged from 11. 4% to78.1%. Ethyl acetate extract was rich in total catechins content (46.6% to 75.9%). Water extracts had significantly (p≤0.05) high amounts of total catechins (12.3% to 27.6%) compared to residual extracts (5.4% to 8.9%) contrary to methylene chloride extracts which had low total catechin contents (1.2% to 4.5%). Caffeine was predominant in methylene chloride tea extracts and ranged from 29.97% to 58.99% pointing to high solvent affinity. Based on this results, ethyl acetate proved to be the ideal solvent for obtaining extract rich in polyphenols, and methylene chloride for isolation of natural caffeine. For high yield of all polyphenols, water was found to be the solvent of choice of tea extracts without regarding the specific class of compound to be isolated. This study confirmed that specialty tea extracts are a promising source of bioactive compounds, suggesting they could be very attractive for use as dietary supplements, cosmetic ingredients and in food industries.

## INTRODUCTION

Tea *Camellia sinensis* is one of the most consumed beverages globally after water. The tea plant has evergreen leaves from which tea beverage is extracted [15]. Two leaves and a bud are most preferably plucked during harvesting. These are known to be highly rich in polyphenols which influences the quality and taste of the tea [23]. Processed tea are classified as either a CTC (Cut, Tear and Curl) or orthodox depending on extent of leaf disruption or rolling and oxidation period, which gives each tea type a characteristic taste and chemical composition [23]. The Crush, Tear, Curl (CTC) teas and specialty teas/orthodox teas (whole leaf) lies primarily in their production processes and resulting characteristics. CTC teas are produced using a set of rollers that Crush, Tear, and Curl the leaves, leading to a fine granular appearance. This process is efficient and yields a tea that infuses quickly, forming a strong, intense flavour, making it ideal for tea bags and mass-market blends such as masala tea, as it provides a full-bodied flavour. Moreover, CTC teas offer consistency, cost-effectiveness, and suitability for large-scale production. However, the mechanical processing can strip away some of the more delicate flavours and aromas, resulting in a less complex cup. In contrast, specialty teas (yellow, oolong, green orthodox, black orthodox and purple teas) involve processing whole leaves through withering, rolling, oxidizing, and drying into different sizes twists and style. This controlled process allows for greater retention of the leaves’ natural flavours and a wider variety of taste profiles. Specialty teas are known for their superior flavour complexity, higher quality, and the potential for unique and varied sensory experiences. However, they tend to be more labour-intensive and their preparation often requires more skill and time. Specialty teas are generally priced higher than CTC teas due to its higher quality, preserving the flavours, aroma and natural oils of the tea leaves, resulting in a tea that is more complex and flavourful as compared to CTC teas. In essence, while CTC teas cater to convenience, specialty teas offer a richer, more diverse drinking experience, appealing to specialist and specialty markets.

Variation of oxidation period produces varied tea types including: green, white, yellow, oolong and black teas. The green teas are non-fermented tea that are processed by immediately pan-firing or steaming freshly picked green tea leaves to prevent oxidation by polyphenol oxidase enzyme. Green teas have abundance of polyphenols such as catechins [5]. Yellow teas, on the other hand, are non-fermented tea with the leaves steamed, sweltered (leaf is piled, then covered and kept at temperatures between 25°-35°C until it turns yellow), rolled and dried using hot air oven to stop oxidation. The tea promotes yellow colour and gives its unique characteristic of pleasant mellow taste. The yellow tea has high amino acid content and is rich in polyphenols [27]. Oolong tea are partially oxidized teas to give characteristic of both black and green tea as the oxidation period is in between these two tea types [13]. The black tea undergoes through multistep processing of withering, rolling, oxidation and drying. Oxidation causes polyphenols to polymerize forming larger and more complex polyphenols: theaflavins (TFs) and thearubigins (TRs) [17], which give the tea type its quality characteristics. It is a product that gives a deep red to dark brown colour liquor and offers delicate sweet and at times a noteworthy astringency taste.

Tea biomolecules consists of amino acid, theanine, alkaloids such as caffeine, theobromine and theophylline, phenolic acids like gallic acid and theogallin, flavonols like quercetin, myricetin and kaempferol and catechins including; epigallocatechin gallate (EGCG), epigallocatechin (EGC),epicatechin gallate (ECG), catechins (C), epicatechin (EC) and gallocatechin gallate (GCG) [11,29,34]. The EGCG has been reported to be responsible for the most potent physiological functions such as prevention of platelet aggregation, promotion of weight loss through fat oxidation, lowering of cholesterol level and inhibition of lipid peroxidation [14,32,43]. The TFs and TRs are characteristic products of quality black tea. Studies have shown that consumption of theanine in tea lowers blood pressure as well as regulates the physiological levels of norepinephrine, dopamine and serotonin levels in the brain[38]. Caffeine can stimulate the central nervous system and cause relaxation of respiratory and cardiac muscles. Phenolic acids, catechins and their derivatives have anti-carcinogenic and anti-mutagenic effects since they act as protective agents of DNA against free radicals, by inactivating carcinogens and inhibiting enzymes involved in pro-carcinogen activation [36]. Moreover, catechins and their derivatives are considered as therapeutic agents on degenerative diseases and brain aging processes as they serve as possible neuroprotective agents in progressive neurodegenerative disorders such as Parkinson’s and Alzheimer’s diseases [28,35].

Overproduction of CTC teas has caused huge volumes of surpluses, and due to the overreliance on export prices of black tea, prices have continued to decline over time, affecting returns to farmers. To sustain the relevance of tea, the tea industry has embarked on rigorous investments to promote value addition and product diversification. Tea extract processing techniques have been adopted to preserve the intake of dietary phytochemicals, leading to an ever-increasing demand for tea extracts and isolated tea bioactive molecules in pharmaceuticals, food-grade additives, and well-marketed cosmetic products [3,33,41,42]. Specialty tea type a product diversification process has previously not known to the Kenyan market, such as green, white, purple, yellow, orthodox processed, and oolong teas, are being popularized.

Tea Camellia sinensis, adhering to the principle of “like dissolves like.” This principle states that polar solvents are more effective at dissolving polar compounds, whereas non-polar solvents better dissolve non-polar compounds. Tea encompasses a diverse array of bioactive constituents, including polyphenols (such as catechins), alkaloids (such as caffeine), amino acids (such as theanine), and various vitamins, each exhibiting distinct polarities. For instance, polyphenols and amino acids are predominantly polar due to their hydroxyl and carboxyl groups, making them highly soluble in polar solvents like water. Conversely, alkaloids like caffeine possess less polarity and are optimally extracted using solvents with low polar solvents such as methylene chloride. Ethyl acetate, characterized by its intermediate polarity, serves as an effective solvent for extracting moderately polar compounds, including caffeine and certain flavonoids. This intermediate polarity of ethyl acetate allows for the selective dissolution of compounds that are not as readily extracted by highly polar or non-polar solvents. The specificity of solvent polarity towards these bioactive compounds ensures an efficient extraction process by maximizing the solubility and stability of the target molecules. This specificity, in turn, significantly influences the antioxidant, anti-inflammatory, and other therapeutic properties of the resultant tea extract. Therefore, the strategic selection of solvents based on their polarity, incorporating polar solvents (water), intermediate polar solvents (ethyl acetate), and less polar solvents (hexane, methylene chloride), is critical for optimizing the yield and purity of specific bioactive compounds from tea leaves. This comprehensive approach facilitates the effective isolation and utilization of tea’s diverse bioactive profile, enhancing its potential health benefits.

Tea extract is an economically attractive source of bioactive constituents, such as polyphenols. Due to the wide variety of phenolic structures, no single solvent or procedure can optimally extract all phenolic compounds from plant materials. Most studies have focused on CTC teas, while specialty teas and the effect of different solvents on their polyphenol content remain understudied. This study aims to identify and quantify the primary bioactive compounds in specialty tea extracts using various solvent treatments.

## MATERIALS AND METHODS

### Materials

#### Chemical and reagents

Gallic standard of phenolic compounds was purchased from Sigma Aldrich Chem (Merck, United States) through Kobian Limited, Nairobi. Standards of catechins were purchased from Sigma Aldrich (Merck, United States) Chem through Kobian limited Nairobi Kenya. (EGC, EGCG, EC, ECG, and +C). The HPLC gradient grade acetonitrile, ethyl acetate, methylene chloride, sodium carbonate, ethylene diamine tetra acetic acid (EDTA), acetic acid, ascorbic acid, Folin-Ciocalteu reagent from Merck (Darmstadt, Germany) were purchased from Sigma Aldrich Chem (Merck, United States) through Kobian Limited Nairobi, Kenya.

### Tea samples

The tea samples for this study were sourced from Tea Research Institute (TRI)-Kericho, situated in the western highlands of Kenya at latitude 0° 22′′ S, longitude 35° 21′′ E and altitude of 2180m above sea level. Fresh two leaves and a bud of green and purple coloured tea leaves were collected from the fields and transported to TRI miniature factory for processing. Japanese green orthodox and matcha tea samples were also included in this study.

### Sample preparation

#### Oolong processing

Fresh tea leaves were sun withered for 30 min before being withered for 1hr indoors. The following step was tossing with a weaved basket for 3-5 minutes, then indoor wither for 2h, tossing again for 3 min, indoor wither for 1hr and finally tossing for 6 min before fixing with a pan fryer to obtain a moisture content of 60%. After that, they were rolled with a roller machine and dried for 1hr at110°C using a microwave oven (Samsung-Model-GE 109 MST). Final product was milled using a coffee miller (Model-Moulinex, Type AR11) and stored in a sealed aluminium bags awaiting analysis.

#### Green processing

Fresh green and purple leaves were indoor withered for 4hr, steamed for 1min, indoor withered again to achieve a moisture content of 60%, rolled with a roller machine, then dried at 70°C using a microwave oven (Samsung-Model-GE 109 MST). The tea was milled with a coffee miller (Model-Moulinex, Type AR11) and stored in sealed aluminum bags until use for analysis.

#### Yellow processing

Fresh green leaves were indoor withered for 4hr, steamed for 6min, followed by indoor withered again to achieve a moisture content of 60%, rolled with a roller machine, and sealed yellowed for 12-18hr, during which the leaves were piled, wrapped in a muslin cloth, and kept damp at 26°C until it turned yellow. They were then rolled using a roller machine, dried with a microwave oven (Samsung-Model-GE 109 MST) at low temperatures of 70°C and milled with a coffee miller (Model-Moulinex, Type AR11) before being stored in sealed aluminum bags awaiting analysis.

#### Black processing

Fresh green tea leaves were withered for 16hr indoors, rolled and oxidized for 3hr. The next phase was drying using a microwave oven (Samsung-Model-GE 109 MST) set at 120°C. For examination, the tea was milled with a coffee miller (Model-Moulinex, Type AR11) and stored in sealed aluminum bags awaiting analysis.

#### Tea samples processing

Tea samples were processed into purple orthodox, green orthodox, yellow orthodox, black orthodox and oolong teas by physical wither, oxidation, rolling and drying according to (manual for tea processing in Kenya). All tea samples were grounded using a coffee grinder and stored in a sealed sterilized aluminium bags for analysis.

### Preparation of tea extract

#### Extraction and fractionation

Sample extraction and fractionation was carried out according to Mariem [25] as shown in Figure 1. Briefly, 50 g of crude grounded tea leaves was extracted with 500 ml of pure distilled water at temperatures of 100^0^C for 1h, under continuous mechanical shaker. The extracts were filtered using Whatman No. 2 filter paper and subjected to separation partition using methylene chloride and ethyl acetate in the ratio of 1:1(v/v). Filtered samples were concentrated using a rotary evaporator and freeze dried.

**FIGURE 1:**
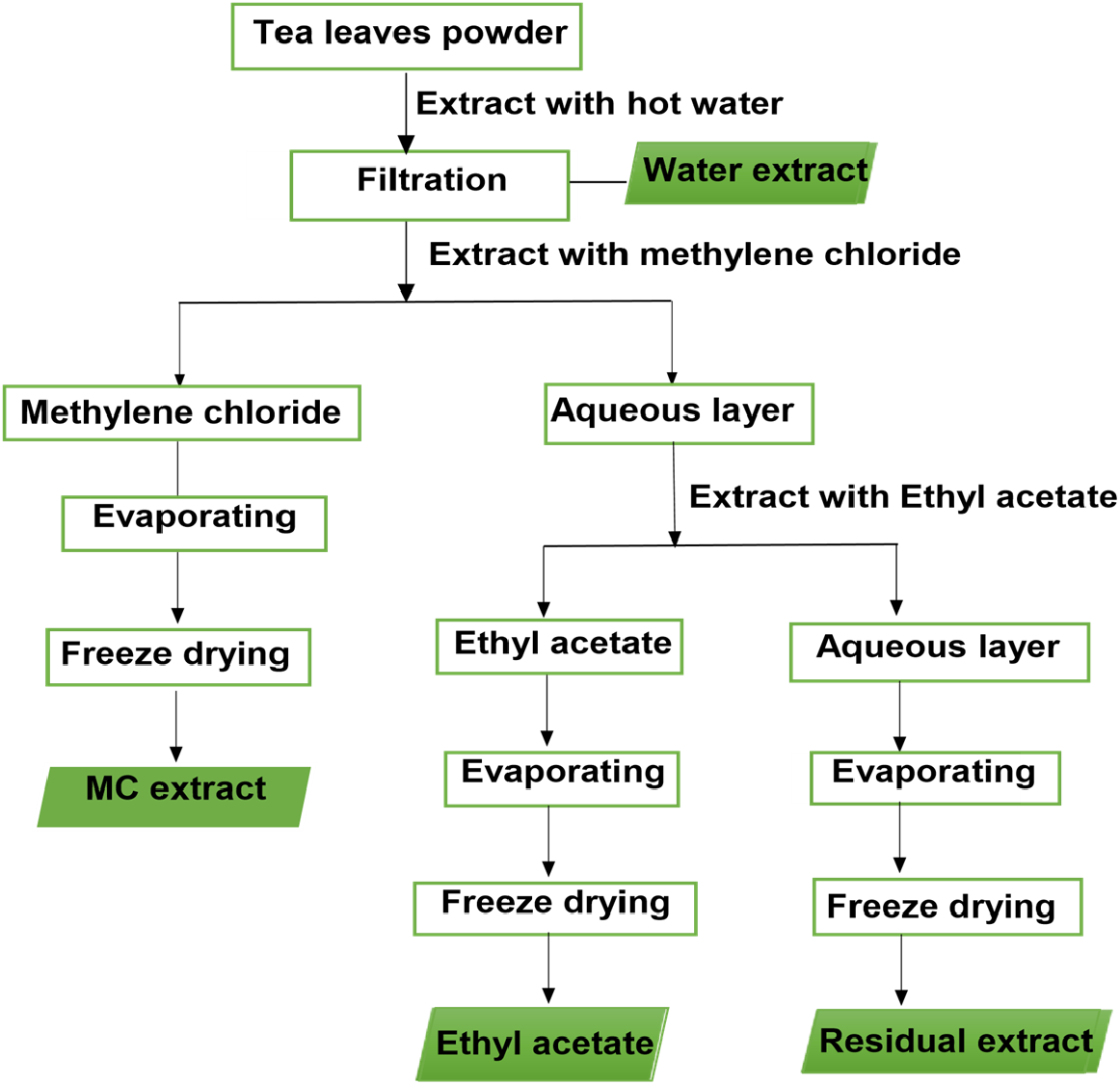
Schematic illustration of the extraction and fractionation procedure [25].

### Determination of extraction yield

Yield weight of each extract based on the dried weight basis was calculated from equation shown below.

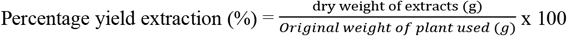

### Determination of total polyphenols

#### Determination of total phenolic content

The total phenolic content of tea extracts was determined according to the spectrophotometric method of International Standards Organization (ISO) ISO-14502-1:2005E. UV/VIS spectrophotometer, Model UV-1800, Shimadzu, Japan was used to determine absorbance at 765nm wavelength [20]. The results were presented as a mean value of triplicate tests.

### Quantitative analysis of selected catechin and caffeine compounds

An International Organization for Standardization for high performance liquid chromatography method (ISO-14502-2:2005E) was used to assay for tea catechins. Determination of the selected compounds was done using Shimadzu, Japan LC 20 AD HPLC system fitted with a SIL 20AC auto sampler, SPD-M20A photo diode array detector [20]. The results were presented as a mean value of triplicate tests.

### Statistical analysis

All statistical analyses were carried out using the SAS version 9.1 package. Data was presented as tables and least significant difference (LSD) values were used to separate differences among the treatment means.

## RESULTS AND DISCUSSION

### Extraction yield

The extraction yields (v/w) of selected specialty teas extracts ranged between 2% to 80% with residual extracts being the highest followed by water, ethyl acetate and methylene chloride being the least as shown in Figure 2. Yield of residual water extracts (82.2%) was significantly higher than that of water extract (28.9%), which was in turn significantly higher than that of ethyl acetate extract (15.1%). However, methylene chloride expressed much lower recovery of polar compounds (2.63%). The comparison between the yields of different fractions showed that content extractable compounds decreased with decreasing solvent polarity [2] which was in agreement with solvents as used in the study; water < ethyl acetate < methylene chloride. Similarly, it has been reported that water generally give a higher total yield of tea extracts compared to ethyl acetate [7,12]. Water has high affinity for extracting considerable amounts of polar bulk compounds while ethyl acetate possesses higher selectivity towards less polar compounds [9], thus lower recovery of polar compounds as observed in this study. In addition, methylene chloride a less polar solvent [1], gives lower recovery of polar compounds, thus the nature of the compound and solvent influences the extraction power. The results from this study therefore provide valuable information on different solvents and their extraction efficiency on specific tea biomolecules which can be exploited in pharmaceutical and food industries given the health benefits of tea.

**FIGURE 2:**
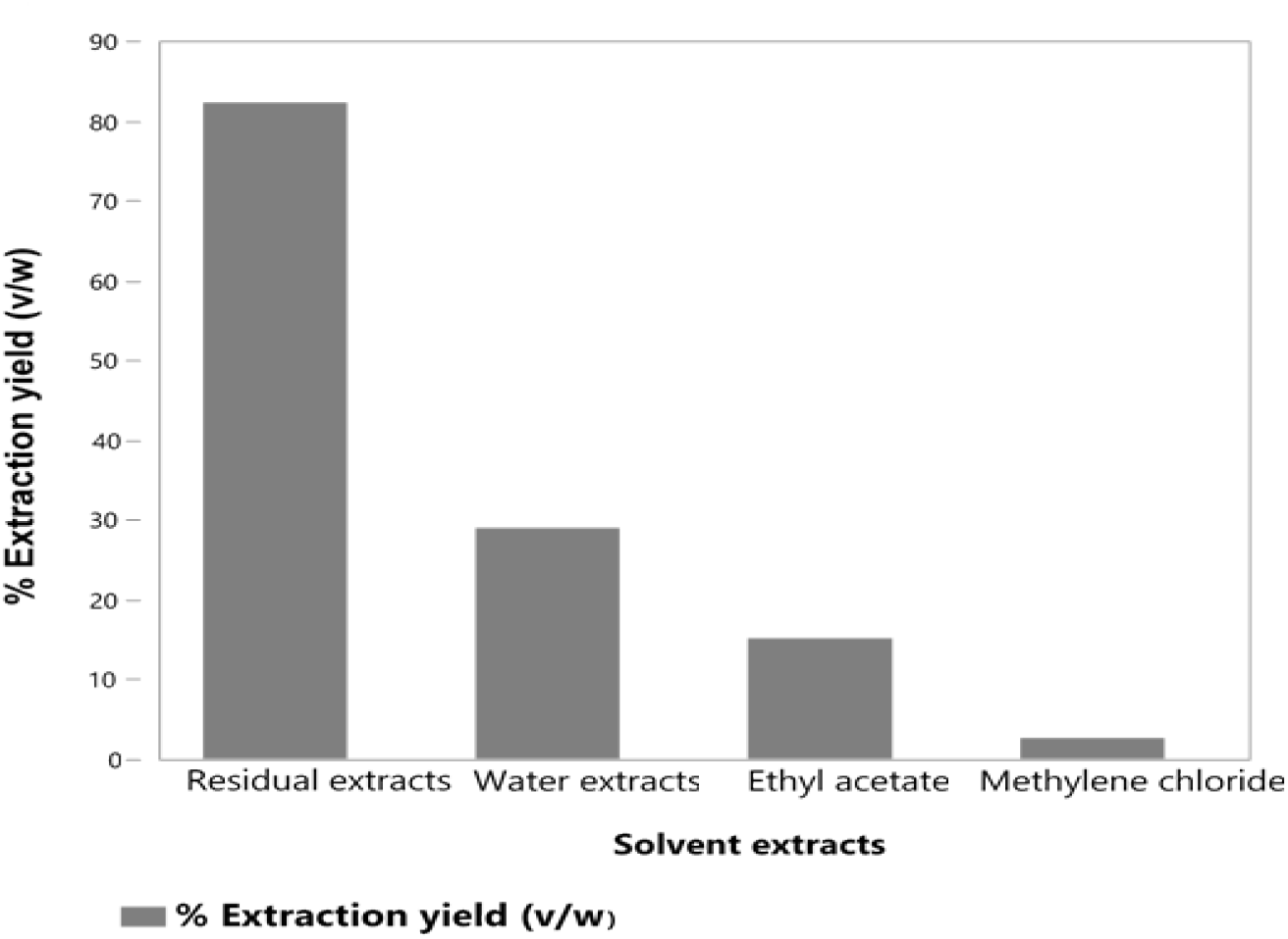
Extraction yields of selected specialty teas in different solvents.

### Total polyphenols

The results showed that total polyphenol content was significantly different (p≤0.05) depending on extraction solvent used and tea type (Table 1). From the results, ethyl acetate extracts were rich in total polyphenols as compared to water, residual and methylene chloride extracts for all tea types. Similar findings on high total polyphenol content in ethyl acetate tea extracts has been reported previously [12]. Previous research findings indicated that water give considerably higher total polyphenols compared to organic solvents [12]. For example, though ethyl acetate possesses higher selectivity towards less polar polyphenols and generates extracts with higher purity,methylene chloride shows low affinity on polar polyphenols [4]. Since polar solvents contain bonds with different electro-negativities, they have large dipole moment (partial charges), which quantify the polarity of solvents[31] as was observed in this study.

**TABLE 1:** Total phenol content (%) and total catechin content (%) of tea extracts used in this study.

| Tea type | Tea extracts | Polyphenol (%) | Total Cat (%) |
| --- | --- | --- | --- |
| Black Orthodox | Water extract | 20.9h | 12.3i |
|  | MC extract | 9.32m | 1.24p |
|  | EA extract | 78.1c | 46.6c |
|  | Residual extract | 12.2k | 5.36l |
|  | Mean | 30.14 | 16.4 |
| Oolong | Water extract | 33.0f | 21.6f |
|  | MC extract | 10.6lm | 4.57m |
|  | EA extract | 83.8b | 56.6b |
|  | Residual extract | 14.2j | 7.69k |
|  | Mean | 35.4 | 22.6 |
| Green orthodox | Water extract | 37.6de | 23.6e |
|  | Orthodox green Japan | 22.8g | 17.8g |
|  | Matcha tea | 11.1kl | 3.73n |
|  | MC extract | 11.2kl | 3.74n |
|  | EA extract | 91.1a | 75.9a |
|  | Residual extract | 18.1i | 8.87j |
|  | Mean | 31.96 | 22.3 |
| Yellow orthodox | Water extract | 38.3d | 27.6d |
|  | MC extract | 11.4kl | 4.50m |
|  | EA extract | 91.9a | 73.9a |
|  | Residual extract | 18.1i | 8.8j |
|  | Mean | 39.9 | 28.7 |
| Purple orthodox | Water extract | 36.2e | 15.9h |
|  | MC extract | 11.3kl | 2.93o |
|  | EA extract | 90.4a | 54.9b |
|  | Residual extract | 17.2i | 5.60l |
|  | Mean | 38.8 | 21.9 |
|  | Total mean | 34.9 | 22.2 |
|  | CV (%) | 2.78 | 1.38 |

MC -Methylene chloride fraction; EA -Ethyl acetate fraction
<sup>ab</sup> Means within a column followed by the same letter are not significantly different at $p \leq 0.05$ according to Duncan's Multiple Range Test (DMRT).

Results from this study corroborated with studies [19,22,26] that non-fermented teas (green orthodox, yellow orthodox and purple orthodox teas) had high total polyphenol content as compared to extracts of semi-fermented (oolong tea) and fermented tea (black orthodox tea). Kenyan teas had high levels of total polyphenols compared to Japanese tea (green orthodox and matcha tea) irrespective of the tea type which was corroborated with previous studies [26,40]. Variation of total polyphenol content between the tea types is attributed by degradation of polyphenols during processing. Polyphenols content in non-fermented teas are high as a result of steaming at the initial stage of non-fermented tea production which inactivates endogenous enzyme polyphenol oxidase involved in the oxidation and hydrolysis of the chemical constituents of the leaves [34]. As for fermented and semi-fermented tea types, the enzymes play important roles during processing. The complex enzymatic reaction of polyphenol oxidase and polyphenolic compounds present in tea leaves leads to formation of theaflavins and thearubigins responsible for the formation of the characteristic colour and flavour of oxidized tea [17].

### Total Catechin levels

Ethyl acetate extracts were rich in total catechins content in all tea samples and ranged between 46.6% to 75.9% (Table 1). Green orthodox EA tea extracts exhibited high levels of total catechins (75.9%). Water extracts had significantly (p≤0.05) high amounts of total catechins as compared with residual extracts contrary to methylene chloride extracts which had low total catechin content (1.24% to 4.57%) of dry weight extract. The data also revealed that, different tea types significantly differed (p≤0.05) in total catechins content with green orthodox tea (75.9%), yellow orthodox tea (73.8%,) oolong tea (56.6%), purple orthodox tea (54.9%) and black orthodox tea (46.6%) extracts as shown in the Table 2. Green orthodox tea extracts from Kenya tea cultivars had significantly (p≤0.05) high total catechins content than those from Japanese green orthodox tea extracts with 23.6% and 17.8%, respectively.

**TABLE 2:**
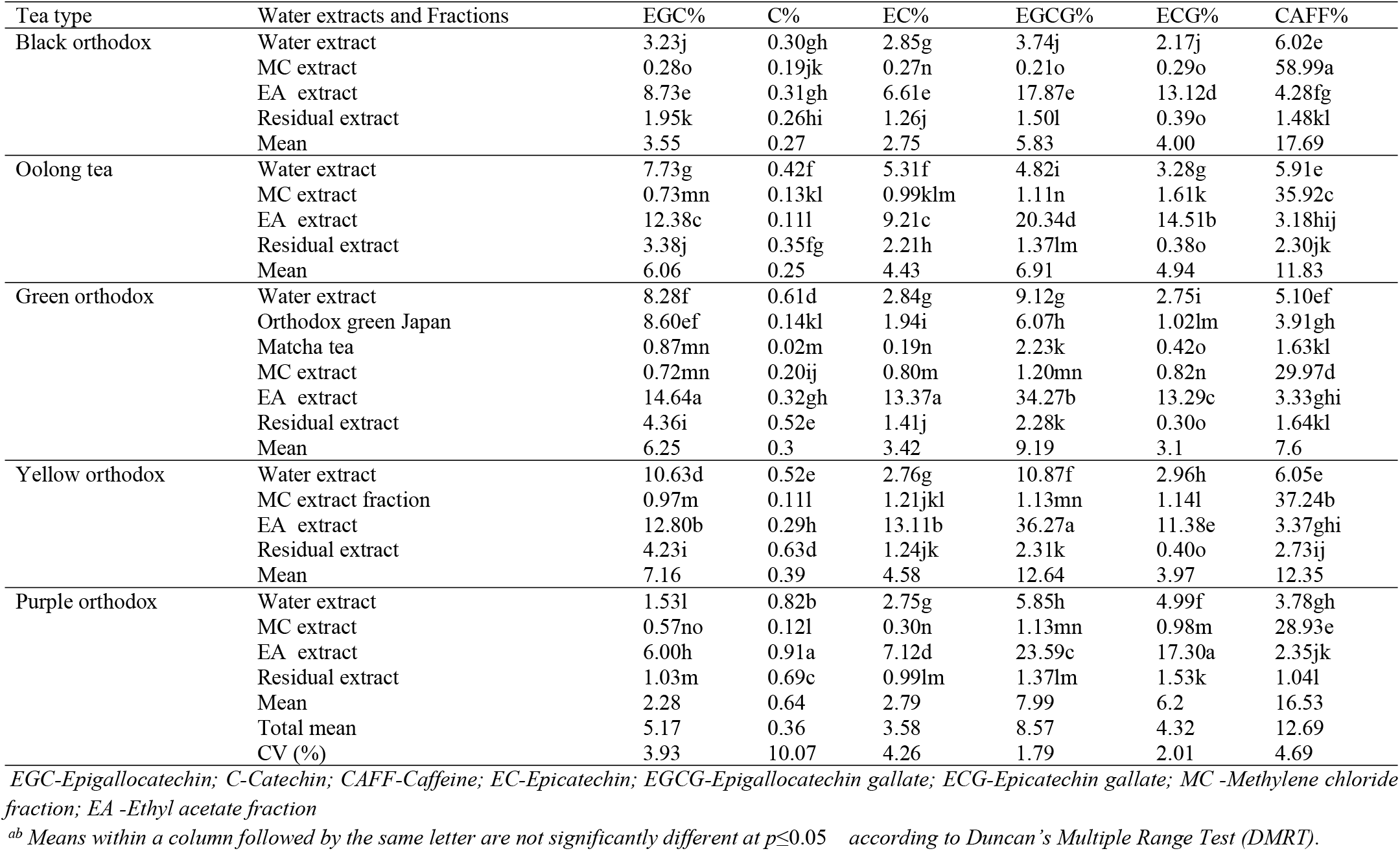
Individual catechins (%) of crude extract and fractions of tea extracts used in this study

| Tea type | Water extracts and Fractions | EGC% | C% | EC% | EGCG% | ECG% | CAFF% |
| --- | --- | --- | --- | --- | --- | --- | --- |
| Black orthodox | Water extract | 3.23j | 0.30gh | 2.85g | 3.74j | 2.17j | 6.02e |
|  | MC extract | 0.28o | 0.19jk | 0.27n | 0.21o | 0.29o | 58.99a |
|  | EA extract | 8.73e | 0.31gh | 6.61e | 17.87e | 13.12d | 4.28fg |
|  | Residual extract | 1.95k | 0.26hi | 1.26j | 1.50l | 0.39o | 1.48kl |
|  | Mean | 3.55 | 0.27 | 2.75 | 5.83 | 4.00 | 17.69 |
| Oolong tea | Water extract | 7.73g | 0.42f | 5.31f | 4.82i | 3.28g | 5.91e |
|  | MC extract | 0.73mn | 0.13kl | 0.99klm | 1.11n | 1.61k | 35.92c |
|  | EA extract | 12.38c | 0.11l | 9.21c | 20.34d | 14.51b | 3.18hij |
|  | Residual extract | 3.38j | 0.35fg | 2.21h | 1.37lm | 0.38o | 2.30jk |
|  | Mean | 6.06 | 0.25 | 4.43 | 6.91 | 4.94 | 11.83 |
| Green orthodox | Water extract | 8.28f | 0.61d | 2.84g | 9.12g | 2.75i | 5.10ef |
|  | Orthodox green Japan | 8.60ef | 0.14kl | 1.94i | 6.07h | 1.02lm | 3.91gh |
|  | Matcha tea | 0.87mn | 0.02m | 0.19n | 2.23k | 0.42o | 1.63kl |
|  | MC extract | 0.72mn | 0.20ij | 0.80m | 1.20mn | 0.82n | 29.97d |
|  | EA extract | 14.64a | 0.32gh | 13.37a | 34.27b | 13.29c | 3.33ghi |
|  | Residual extract | 4.36i | 0.52e | 1.41j | 2.28k | 0.30o | 1.64kl |
|  | Mean | 6.25 | 0.3 | 3.42 | 9.19 | 3.1 | 7.6 |
| Yellow orthodox | Water extract | 10.63d | 0.52e | 2.76g | 10.87f | 2.96h | 6.05e |
|  | MC extract fraction | 0.97m | 0.11l | 1.21jkl | 1.13mn | 1.14l | 37.24b |
|  | EA extract | 12.80b | 0.29h | 13.11b | 36.27a | 11.38e | 3.37ghi |
|  | Residual extract | 4.23i | 0.63d | 1.24jk | 2.31k | 0.40o | 2.73ij |
|  | Mean | 7.16 | 0.39 | 4.58 | 12.64 | 3.97 | 12.35 |
| Purple orthodox | Water extract | 1.53l | 0.82b | 2.75g | 5.85h | 4.99f | 3.78gh |
|  | MC extract | 0.57no | 0.12l | 0.30n | 1.13mn | 0.98m | 28.93e |
|  | EA extract | 6.00h | 0.91a | 7.12d | 23.59c | 17.30a | 2.35jk |
|  | Residual extract | 1.03m | 0.69c | 0.99lm | 1.37lm | 1.53k | 1.04l |
|  | Mean | 2.28 | 0.64 | 2.79 | 7.99 | 6.2 | 16.53 |
|  | Total mean | 5.17 | 0.36 | 3.58 | 8.57 | 4.32 | 12.69 |
|  | CV (%) | 3.93 | 10.07 | 4.26 | 1.79 | 2.01 | 4.69 |

EGC-Epigallocatechin; C-Catechin; CAFF-Caffeine; EC-Epicatechin; EGCG-Epigallocatechin gallate; ECG-Epicatechin gallate; MC -Methylene chloride fraction; EA -Ethyl acetate fraction
<sup>ab</sup> Means within a column followed by the same letter are not significantly different at $p \leq 0.05$ according to Duncan's Multiple Range Test (DMRT).

### Individual catechins and caffeine levels

In order to fully elucidate a relation between the type of solvent and the amount of bioactive compounds extracted, quantitative analysis was conducted by HPLC–PDA (Figure 3). The compounds profiled were mainly individual catechins, also known as flavanols and caffeine. Yellow, green and purple orthodox teas had significantly (p≤0.05) high content of individual catechins compared to oolong and black orthodox teas (Table 2). Results indicated that catechins, EGCG, EGC, EC and ECG were the most abundant compounds in all ethyl acetate tea extracts (Table 2). Catechins forms a hydrogen bonding through the dipole-dipole interaction that results from the polarity of catechins and water [31]. However, catechin has a great affinity for ethyl acetate that results from greater attraction of organic solvent ethyl acetate and organic compound catechins, “like dissolves like” [10]compared to water an inorganic solvent. In addition, ethyl acetate has a low dipole moment (partial charges) and small dielectric constant (indicates how easily a material can become polarized by imposition of an electric field on an insulator) than water thus a less polar solvent and highly selectivity to less polar flavanols and produce of pure extracts. Ethyl acetate was confirmed to be an appropriate solvent for isolating catechins from tea extract, based on concentration of catechins.

**FIGURE 3:**
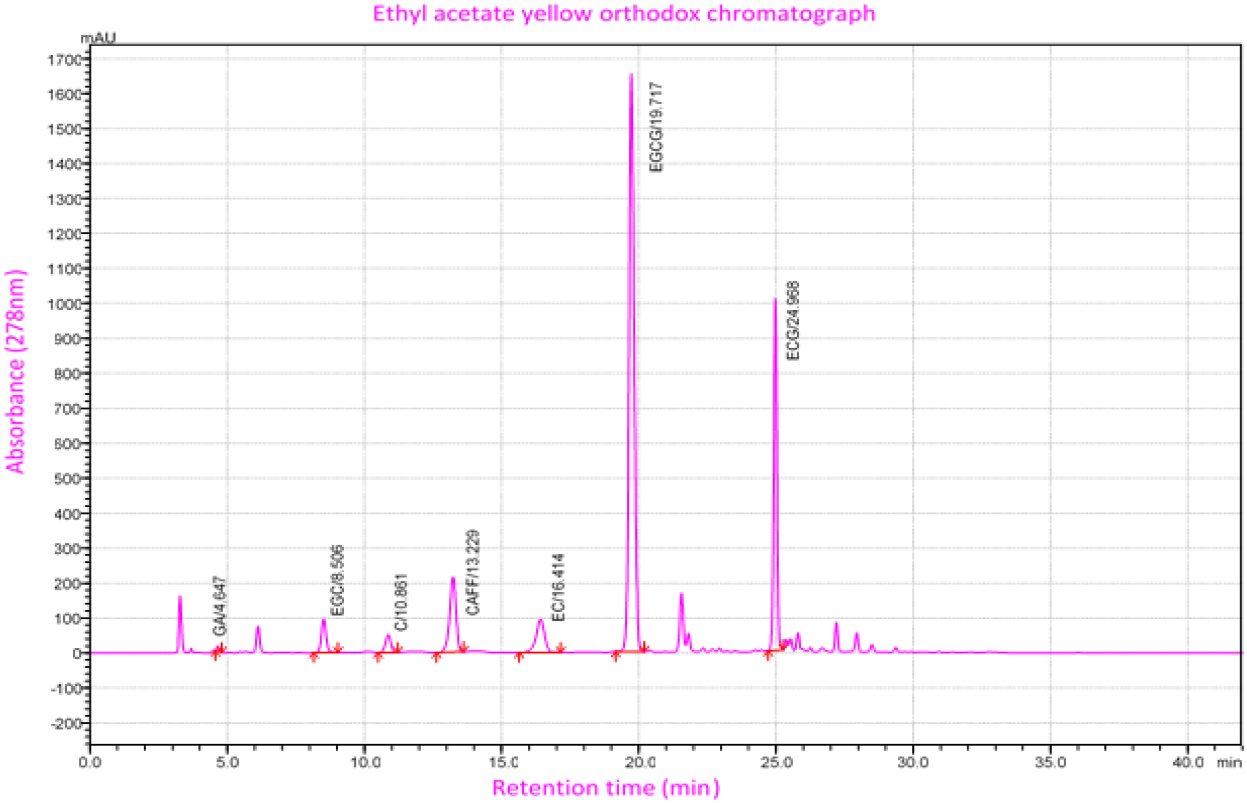
Ethyl acetate yellow orthodox chromatograph EGC-Epigallocatechin; C-Catechin; CAFF-Caffeine; EC-Epicatechin; EGCG-Epigallocatechin gallate; ECG-Epicatechin gallate

Caffeine was significantly (p≤0.05) high in black orthodox tea compared to all mentioned tea types and predominant biomolecule in methylene chloride tea extract (Table 2). Caffeine has a stronger attraction to water due to the dipole-dipole interaction that results from the greater polarity of caffeine and the hydrogen bonds that form between caffeine and water [31]. However, caffeine has a greater affinity for methylene chloride and will easily dissolve in this solvent because both caffeine and methylene chloride are organic substances while water is inorganic, thus in methylene chloride, caffeine will have a greater attraction for the organic solvent and the hydrogen bonds between caffeine and water will be broken [39]. For assessing the efficacy of solvents to extract a specific compound, the first requirement was that the solvent efficiently extracts the desired compound from all tested specialty tea extracts. This is important because food industry is more focused on the isolation of food colorants or preservatives, while the pharmaceutical industry needs biologically active compounds.

In terms of different oxidation period, total catechins content varied significantly (p≤0.05) among the different tea types. Non-aerated teas had high total catechins content than semi-aerated and aerated teas. However, purple orthodox extracts, rich in polyphenols known as anthocyanins, that gives the purple leaf its purple-pigment, exhibited low levels of catechins content compared to green orthodox extracts. The extent of variation of total catechins levels is as a result of predominant anthocyanins in purple leaf which is not detectable at 278nm wavelength used for catechins analysis.

The extraction yield depends on many factors, where the most important are the type and polarity of the solvent applied. However, there is no unique solvent or extraction procedure optimal for obtaining all polyphenols from plant material such as tea [9].Water has been widely used for the extraction of[16,37] phenolic compounds from tea, because it is suitable, nontoxic for researchers and the environment (cite). Many studies have been published on its usage as a solvent for extraction of phenolics [21,26,30] but compared to other solvents in our study, water did not show the best extracting ability. Several studies have indicated methylene chloride as a solvent of choice for caffeine extraction from tea [18,20,39]. In one study, combination of ethyl acetate and dichloromethane were evaluated, and ethyl acetate expressed the highest selectivity towards catechins compounds[8], which is similar to our findings. By possessing a low boiling point and toxicity ethyl acetate could safely be used in food, cosmetic and pharmaceutical industry for extracting polyphenols from tea. Additionally, ethyl acetate is also good from economical point of view as it is inexpensive [6,24].The results obtained herein support its usage in specialty tea extract utilization as ethyl acetate proved to be the most efficient solvent for extracting investigated polyphenols.

## CONCLUSION

Choosing an optimal solvent for extraction of beneficial natural products from specialty teas extract is dependent on many factors among them the bioactive compounds to be obtained. Methylene chloride was established as the most applicable optimal solvent for extraction of natural caffeine from all tea types. Hot water presents the best option for industries that target to achieve a high yield of polyphenols without regarding the purity of the compounds extracted. This study gives useful and valuable information for efficient exploitation of specialty teas in industry. These tea extracts are a rich source of polyphenols, making them valuable for various industries, including pharmaceutical, cosmetic, and food sectors. Utilizing these extracts can also boost the economy by creating new market opportunities and promote ecological benefits by encouraging sustainable practices.

## Acknowledge

This work was supported by Tea Research Institute-(TRI-Kericho), Kenya Medical Research Institute-(KEMRI-Nairobi), University of Kabianga –Kericho and Tumoi tea Limited-Nandi hills.

## REFERENCES

[1] Aljamali, N.M. (2016). Physical Properties of liquids. Polar 17(1.8), 0–9.

[2] Altemimi, A.; Lakhssassi, N.; Baharlouei, A.; Watson, D.G.; Lightfoot, D.A. (2017). Phytochemicals: Extraction, isolation, and identification of bioactive compounds from plant extracts. Plants 6(4), 42.

[3] Badria, F.A.; Zidan, O.A. (2004). Natural products for dental caries prevention. J. Med. Food 7(3), 381–4.

[4] Banerjee, S.; Chatterjee, J. (2015). Efficient extraction strategies of tea (Camellia sinensis) biomolecules. J. Food Sci. Technol. 52(6), 3158–3168

[5] Bieleski, V.M. (1994). Willson, KC and Clifford. MN (editors). Tea. Cultivation to consumption. Chapman & Hall, London: 1992. Pp xx, 769

[6] Bonilla, F.; Mayen, M.; Merida, J.; Medina, M. (1999). Extraction of phenolic compounds from red grape marc for use as food lipid antioxidants. Food Chem. 66(2), 209–15.

[7] Chan, E.W.C.; Soh, E.Y.; Tie, P.P.; Law, Y.P. (2011). Antioxidant and antibacterial properties of green, black, and herbal teas of Camellia sinensis. Pharmacognosy Res. 3(4), 266.

[8] Choung, M.; Hwang, Y.; Lee, M.; Lee, J.; Kang, S.; Jun, T. (2014). Comparison of extraction and isolation efficiency of catechins and caffeine from green tea leaves using different solvent systems. Int. J. Food Sci. Technol. 49(6), 1572–8.

[9] Dai, J.; Mumper, R.J. (2010). Plant phenolics: extraction, analysis and their antioxidant and anticancer properties. Molecules 15(10), 7313–52.

[10] Dong, J.-J.; Ye, J.-H.; Lu, J.-L.; Zheng, X.-Q.; Liang, Y.-R. (2011) Isolation of antioxidant catechins from green tea and its decaffeination. Food Bioprod. Process. 89(1), 62–6.

[11] Enzveiler, L.; Gressler, G.; Heckler, E.; Picoli, S.; Suyenaga, E.S. (2011). Evaluation of antimicrobial activity of aqueous extract of white tea Camellia sinensis L. Kuntze (1887). Pharmacol. 2(5), 2011.

[12] Erol, N.T.; Sari, F.; Polat, G.; Velioglu, Y.S. (2009). Antioxidant and antibacterial activities of various extracts and fractions of fresh tea leaves and green tea. Tarim Bilim. Derg. 15(4), 371– 8.

[13] Fanaro, G.B.; Duarte, R.C.; Santillo, A.G.; e Silva, M.E.M.P.; Purgatto, E.; Villavicencio, A. (2012). Evaluation of γ-radiation on oolong tea odor volatiles. Radiat. Phys. Chem. 81(8), 1152– 6.

[14] Hara, Y. (2011) Tea catechins and their applications as supplements and pharmaceutics. Pharmacol. Res. 64(2), 100–104.

[15] Hazra, A.; Saha, J.; Dasgupta, N.; Sengupta, C.; Kumar, P.M.; Das, S. (2017). Health-benefit assets of different indian processed teas: a comparative approach. Am. J. Plant Sci. 8(07), 1607.

[16] Huang, W.-Y.; Lin, Y.-R.; Ho, R.-F.; Liu, H.-Y.; Lin, Y.-S. (2013). Effects of water solutions on extracting green tea leaves. Sci. World J. 2013.

[17] Imran, A.; Arshad, M.U.; Mehmood, S.; Ahmed, R.S.; Butt, M.S.; Ahmed, A.; Imran, M.; Arshad, M.S.; Faiza, N.; Haq, I. (2018). Oxidative Stress Diminishing Perspectives of Green and Black Tea Polyphenols: A Mechanistic Approach. Polyphenols, 25.

[18] Jara-Palacios, M.J.; Hernanz, D.; Escudero-Gilete, M.L.; Heredia, F.J. (2014). Antioxidant potential of white grape pomaces: Phenolic composition and antioxidant capacity measured by spectrophotometric and cyclic voltammetry methods. Food Res. Int. 66, 150–7.

[19] Kerio, L.C.; Bend, J.R.; Wachira, F.N.; Wanyoko, J.K.; Rotich, M.K. (2011). Attenuation of t-Butylhydroperoxide induced oxidative stress in HEK 293 WT cells by tea catechins and anthocyanins. J. Toxicol Env. Heal. Sci 3, 367–75.

[20] Kim, W.-J.; Kim, J.-D.; Kim, J.; Oh, S.-G.; Lee, Y.-W. (2008). Selective caffeine removal from green tea using supercritical carbon dioxide extraction. J. Food Eng. 89(3), 303–9.

[21] Koech, K.R.; Wachira, F.N.; Ngure, R.M.; Wanyoko, J.K.; Bii, C.; Karori, S.M. (2013). Antibacterial and synergistic activity of different tea crude extracts against antibiotic resistant S. aureus, E. coli and a clinical isolate of S. typhi. Sci. J. Microbiol. 3: 2276–626X.

[22] Koech, R.; Wachira, F.; Ngure, R.; Wanyoko, J.; Bii, C.; Karori, S.; Kerio, L.; Koech, K.; Wachira, F.; Ngure, R. (2013). Antimicrobial, synergistic and antioxidant activities of tea polyphenols. Microb. Pathog. Strateg. Combat. Them 2, 971–81.

[23] Kosińska, A.; Andlauer, W. (2014). Antioxidant capacity of tea: Effect of processing and storage. Processing and impact on antioxidants in beverages, Elsevier, pp. 109–20.

[24] Louli, V.; Ragoussis, N.; Magoulas, K. (2004). Recovery of phenolic antioxidants from wine industry by-products. Bioresour. Technol. 92(2), 201–8.

[25] Mariem, S.; Hanen, F.; Inès, J.; Mejdi, S.; Riadh, K. (2014). Phenolic profile, biological activities and fraction analysis of the medicinal halophyte Retama raetam. South African J. Bot. 94, 114–21.

[26] Mbuthia, S.K.; Wachira, F.N.; Koech, R.K. (2014). In-vitro antimicrobial and synergistic properties of water soluble green and black tea extracts. African J. Microbiol. Res. 8(14), 1527– 34.

[27] Ning, J.; Li, D.; Luo, X.; Ding, D.; Song, Y.; Zhang, Z.; Wan, X. (2016). Stepwise identification of six tea (Camellia sinensis (L.)) categories based on catechins, caffeine, and theanine contents combined with fisher discriminant analysis. Food Anal. Methods 9(11), 3242–50.

[28] Ozcan, T.; Akpinar-Bayizit, A.; Yilmaz-Ersan, L.; Delikanli, B. (2014). Phenolics in human health. Int. J. Chem. Eng. Appl. 5(5), 393.

[29] Padmini, E.; Valarmathi, A.; Rani, M.U. (2010). Comparative analysis of chemical composition and antibacterial activities of Mentha spicata and Camellia sinensis. Asian J. Exp. Biol. Sci 1(4), 772–81.

[30] Rashid, K.; Wachira, F.N.; Ngure, R.M.; Nyabuga, J.N.; Wanyonyi, B.; Murilla, G.; Isaac, A.O. (2014). Kenyan purple tea anthocyanins ability to cross the blood brain barrier reinforcing brain antioxidant capacity in mice. African Crop Sci. J. 22, 819–28.

[31] Reichardt, C.; Welton, T. (2011). Solvents and solvent effects in organic chemistry. John Wiley & Sons.

[32] Reygaert, W.C. (2018). Green tea catechins: Their use in treating and preventing infectious diseases. Biomed Res. Int. 9105261-9105261.

[33] Senanayake, S.P.J.N. (2013). Green tea extract: Chemistry, antioxidant properties and food applications–A review. J. Funct. Foods 5(4), 1529–41.

[34] Shahidi, F.; Naczk, M. (2003). Phenolics in food and nutraceuticals. CRC press.

[35] Sharangi, A.B. (2009). Medicinal and therapeutic potentialities of tea (Camellia sinensis L.)–A review. Food Res. Int. 42(5–6), 529–35.

[36] Singh, B.N.; Prateeksha Rawat, A.K.S.; Bhagat, R.M.; Singh, B.R. (2017) .Black tea: Phytochemicals, cancer chemoprevention, and clinical studies. Crit. Rev. Food Sci. Nutr. 57(7), 1394–410.

[37] Tahir, A.; Moeen, R. (2011) .Comparison of antibacterial activity of water and ethanol extracts of Camellia sinensis (L.) Kuntze against dental caries and detection of antibacterial components. J. Med. Plants Res. 5(18), 4504–10.

[38] Too, J.; Wanyoko, J.; Kinyanjui, T.; Moseti, K.; Wachira, F. (2016) .Quantitative Estimation of γ-Glutamylethylamide in Commercially Available Made Teas [Camellia sinensis (L.) O. Kuntze, Theaceae] in Kenya. Am. J. Plant Sci. 7(01), 55.

[39] Vuong, Q.V; Roach, P.D. (2014). Caffeine in green tea: its removal and isolation. Sep. Purif. Rev. 43(2), 155–74.

[40] Wachira, F.N.; Kamunya, S.M. (2005). Kenyan teas are rich in antioxidants. Tea 26(2), 81–9.

[41] Wang, H.; Provan, G.J.; Helliwell, K. (2000) Tea flavonoids: their functions, utilisation and analysis. Trends Food Sci. Technol. 11(4–5), 152–60.

[42] Yilmaz, Y. (2006) Novel uses of catechins in foods. Trends Food Sci. Technol. 17(2), 64–71.

[43] Zaveri, N.N.T. (2006) .Green tea and its polyphenolic catechins: medicinal uses in cancer and noncancer applications. Life Sci. 78(18), 2073–80.

